# CAMP: Coreset Accelerated Metacell Partitioning enables scalable analysis of single-cell data

**DOI:** 10.64898/2025.12.11.693725

**Authors:** Danrong Li, Young Kun Ko, Stefan Canzar

## Abstract

Scaling metacell inference to atlas-level single-cell datasets demands algorithms that are both computationally efficient and geometrically faithful. We introduce CAMP (Coreset Accelerated Metacell Partitioning), a metacell framework that preserves the intrinsic structure of the cellular manifold while enabling scalable analysis of millions of cells. CAMP leverages coreset-based sampling to construct a small, weighted subset of representative cells that approximates the full dataset with provable geometric guarantees. This formulation transforms metacell construction into a coreset inference problem, reducing runtime and memory complexity by up to an order of magnitude without loss of accuracy.

Through extensive experiments, we show that CAMP produces metacells that are compact, well-separated, and biologically coherent, achieving performance on par with or exceeding existing methods including MetaCell, SuperCell, SEACells, and MetaQ. By combining theoretical efficiency with empirical robustness, CAMP establishes coreset acceleration as a principled foundation for scalable, high-fidelity metacell inference in single-cell transcriptomics.

## 1 Introduction

Single-cell RNA sequencing (scRNA-seq) technologies now profile hundreds of thousands of cells in a single experiment, offering an unprecedented view of cellular heterogeneity during development, immunity, and disease [1][2][3][4]. Yet this level of resolution introduces substantial redundancy and noise [5]: many cells exhibit near-identical expression profiles with high sparsity [6][7], while technical variability obscures underlying biological structure. Analyses that operate directly on the single-cell matrix therefore face severe statistical and computational challenges [6][8].

To address these challenges without introducing excessive smoothing [9][10] or generating false positives [11][7], which are known issues of imputation based approaches, a common strategy is to use metacell frameworks [10]. These methods summarize transcriptionally similar cells into coherent groups that represent stable and biologically meaningful expression states, typically obtained by averaging their expression profiles. Over the past few years, several computational strategies have emerged to implement this idea. MetaCell [10] partitions single-cell data into metacells by constructing a balanced k-nearest neighbor (kNN) graph and iteratively refining densely connected subgraphs through resampling, outlier detection, and rebalancing to ensure homogeneous clusters. MetaCell2 [12] extends this approach with a divide-and-conquer strategy that partitions large datasets into manageable subsets and replaces stochastic resampling with a deterministic stability score, improving scalability and sensitivity to rare cell types. SuperCell [5] is an unsupervised framework that constructs metacells by iteratively merging highly similar cells in a kNN graph built using Euclidean distance. It models the data as a graph in which each cell is represented as a singleton node, and then applies the walktrap community detection algorithm to progressively merge densely connected nodes until the desired number of metacells is reached. SEACells [13] begins by constructing a kNN graph using Euclidean distance and converting it into an adaptive Gaussian kernel, which serves as a similarity matrix in a high-dimensional feature space. Archetypal analysis is then performed on this kernel to identify representative “extreme” transcriptional states, where the number of archetypes corresponds to the desired number of metacells. MetaQ [8] formulates metacell construction as a vector quantization problem over single-cell expression profiles, using a deep autoencoder with an encoder, a learnable codebook, and a decoder that reconstructs gene expression to guide the quantization.

Despite their success, existing frameworks share fundamental limitations. Most rely on graph or manifold formulations that require constructing large matrices and performing iterative optimization, leading to beyond superlinear computational cost. They are also sensitive to hyperparameter choices and often depend heavily on GPU resources to remain practical at scale. As single-cell datasets continue to grow into the hundreds of thousands and millions of cells, there is an increasing need for methods that achieve both computational scalability and geometric fidelity without requiring specialized hardware. Even after we applied substantial implementation-level optimizations, SEACells continued to produce out-of-memory errors on large inputs, while MetaCell and MetaCell2 exceeded a 48-hour runtime limit when dataset sizes increased. Independent evaluations in [8] confirm similar behavior, with SEACells and MetaCell2 stalling or failing on datasets in the 100,000 to 200,000 cell range. These limitations collectively make existing approaches impractical for performing metacell partitioning on modern large-scale single-cell data.

In order to make metacell partition feasible for large scale datasets, we introduce CAMP (Coreset Accelerated Metacell Partitioning), a framework designed to meet this need. CAMP builds on the archetypal formulation introduced by SEACells but replaces its iterative Frank–Wolfe optimization with a lightweight geometric coreset [14], through which metacells are inferred according to positional proximity. It yields results that are comparable in metacell quality metrics (see Section 3.2) while drastically reducing computational cost. By anchoring analysis on representative subsets rather than the full data matrix, CAMP bridges theoretical ideas from geometric summarization [14] with practical demands in single-cell transcriptomics, providing an efficient and conceptually straightforward foundation for large-scale metacell inference.

## 2 Methods

CAMP builds on the principles of archetypal analysis [15][16], a convex-hull formulation that identifies a set of extremal points (“archetypes”) capturing the boundary of the cellular state space. These archetypes delineate distinct transcriptional states within the manifold and therefore serve as natural centers for metacell formation (Supplementary Figure 1). A key theoretical insight enabling CAMP is Proposition 1 of [17], which shows that a coreset constructed for the *k*-means objective is also a valid coreset for archetypal analysis, establishing a direct connection between clustering and archetypal inference.

Classical archetypal analysis, including the kernel formulation used in SEACells [13], typically refines archetypes through iterative optimization over the full dataset or its kernel representation, a procedure that becomes computationally prohibitive at atlas scale. CAMP circumvents this bottleneck by selecting a lightweight *k*-means coreset [14] of size *k* when inferring *k* metacells and treating the resulting representative points as archetypes directly. Each cell is then assigned to the archetype that best represents it in the chosen geometric or kernel space in a one-shot setting. This bypasses iterative archetypal refinement entirely while retaining the desirable properties of *k*-means, which is to minimize within-cluster variance and maximize separation, aligning naturally with biological notions of metacell compactness and distinctness. Moreover, *k*means has been extensively studied from a theoretical perspective, particularly through coreset constructions [18][19][20][21][22][23], providing a well-established foundation for scalable and geometry-preserving metacell inference. The specific coreset construction and sampling distribution used in CAMP follow the framework of [14], which we describe in the next section.

### 2.1 Weighted distribution for sampling CAMP coresets

We define the gene expression matrix as 𝒳 ∈ ℝ^*n×d*^, where *n* denotes the number of cells and *d* the number of features (e.g., genes). Each cell is represented as a row vector 𝒳_*i*_ ∈ ℝ^1*×d*^ for *i* ∈ [*n*]. Let *µ* denote the global mean of all cell vectors, and let *d*_*i*_ =∥ 𝒳_*i*_ −*µ* ∥_2_ be the Euclidean distance between the cell *i* and the mean. CAMP selects a subset of representative cells, termed coreset or archetypes, according to the following sampling distribution [14]:

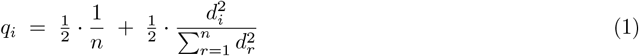

We now introduce the formal notion of a lightweight coreset, which is a small weighted subset approximating the full dataset for solving *k*-means with small relative error.

#### Definition 1

([14, Definition 1. (*ε, k*)-lightweight coreset for *k*-means]). *Let ε* ∈ (0, 1) *and k* ∈ ℕ. *Let* 𝒳⊂ ℝ^*d*^ *be a set of points with mean µ*(𝒳 ). *A weighted set C is an* (*ε, k*)*-lightweight coreset of* 𝒳 *if for every set Q* ⊂ ℝ^*d*^ *of cardinality at most k, the k-means costs on* 𝒳 *and C are within a relative error ε:*

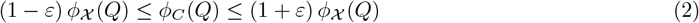

The following theorem shows that sampling points according to (1) produces such a coreset as described in Definition 1 with high probability.

#### Theorem 1

([14, Theorem 2.]). *Let ε* ∈ (0, 1), *δ* ∈ (0, 1) *and k* ∈ ℕ. *Let* 𝒳 *be a set of points in* ℝ^*d*^ *and let coreset C contain cells sampled according to distribution (1) with a sample size m of at least*

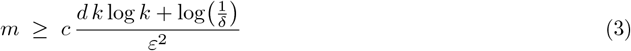

*where c is an absolute constant. Then, with probability all but δ, C is a* (*ε, k*)*-lightweight coreset (see Definition 1) of* 𝒳 *for solving k-means problems*.

### Theoretical Guarantee

Informally, Theorem 1 shows that if the coreset size *m* satisfies

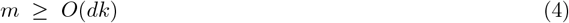

then, with probability all but *δ*, the sampled set *C* forms a (*ε, k*)-lightweight coreset of 𝒳 for the *k*-means objective as described in Definition 1. In other words, the clustering error computed on the coreset *C* approximates that of the full dataset 𝒳 within a multiplicative factor of 1 ± *ε* as stated in (2).

In our setting, the feature dimension *d* corresponds to the number of highly variable genes (HVGs) selected during preprocessing, typically set to the top 2,000 most variable genes. Since *d* ≪ *n* in large single-cell datasets, the required coreset size scales approximately as *m* = *O*(*k*). Therefore, to infer *k* metacells, it suffices to sample on the order of *k* representative cells.

### 2.2 Kernel-based algorithm variants

We introduce CAMP, a family of coreset-accelerated metacell algorithms that share a common lightweight sampling strategy (see Section 2.1) but differ in how distances or similarities are utilized during metacell assignment. CAMP1 serves as our default variant and performs assignment directly in the gene-expression space using Euclidean distances. Motivated by observations in [13] that archetypal formulations tend to emphasize boundary cells while under-representing dense interior regions, we develop CAMP2 and CAMP3 as kernel-based extensions that can capture richer geometric structures. CAMP4 further combines gene-space sampling with kernel-space assignment to improve robustness across heterogeneous transcriptomic landscapes. While all variants follow the same coreset-based sampling pipeline, they exhibit different performance tradeoffs, which form the basis for a practical selection guideline that we introduce in Section 4.

#### Algorithm 1

CAMP1-4

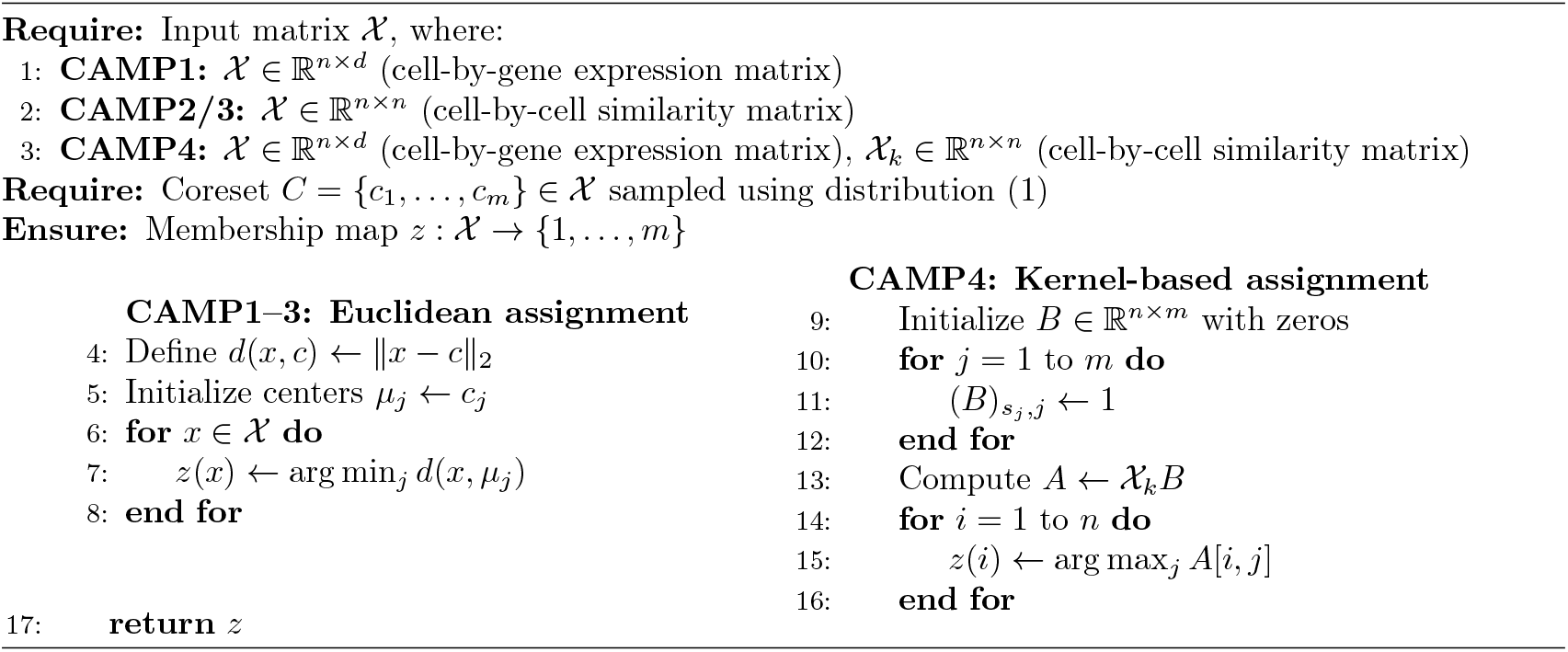

According to Algorithm 1, CAMP1 samples a lightweight geometric coreset and treats the selected points as archetypes. Each remaining cell is then assigned to its nearest archetype in a single step based on Euclidean distance. Optional refinement with a few Lloyd iterations can be applied to adjust cluster boundaries (Supplementary Algorithm 1). Supplementary Figure 1a shows that CAMP1’s coreset provides broad and well-distributed coverage of the transcriptomic manifold, indicating that Euclidean geometry alone is often sufficient to represent the global structure of scRNA-seq data. Supplementary Figure 1b and Figure 1c further confirm that CAMP2 and CAMP3 generate coresets with comparable coverage despite operating in kernel-defined similarity spaces.

To capture higher-order structure that may not be directly reflected in Euclidean distances, CAMP2 replaces raw geometry with similarities derived from the linear kernel

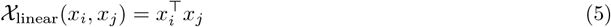

which highlights direction-based relationships between expression vectors and can better distinguish cell states that differ along correlated gene-expression patterns. This representation is particularly effective in high-dimensional spaces, where angular information frequently provides a more stable notion of similarity than pointwise Euclidean distances.

CAMP3 generalizes this idea by incorporating local density information through an adaptive Gaussian kernel [13]. Here, similarities decay with Euclidean distance but are additionally scaled by bandwidth parameters *σ*_*i*_ and *σ*_*j*_ that reflect local neighborhood variability:

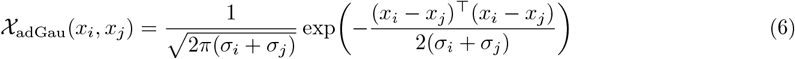

The adaptive term (*σ*_*i*_ + *σ*_*j*_) allows the kernel to expand in sparse regions and contract in dense ones, yielding a similarity landscape that is sensitive to varying transcriptional densities and more robust to dropout or sparsity. This design mirrors the adaptive kernels used in SEACells [13].

Finally, CAMP4 integrates geometric and kernel-based representations in a hybrid strategy. Coreset sampling is first performed in the original gene-expression space to ensure broad geometric coverage, leveraging the efficiency and stability of the Euclidean coreset. Each cell is then assigned to the coreset representative with which it has the highest similarity under the adaptive Gaussian kernel defined in (6), allowing the assignment step to capture nonlinear or density-dependent similarity patterns that Euclidean distance may miss. This two-stage design keeps CAMP4 computationally lightweight while enabling more flexible modeling of cell-cell similarity.

## 3 Results

We used two datasets to benchmark the performance of CAMP against state-of-the-art metacell partitioning approaches, evaluating both metacell quality and downstream performance in a cell-type classification task.

### 3.1 Datasets and benchmarking setup

The first is a single-cell RNA-seq dataset [24] of human peripheral blood mononuclear cells (PBMC) of seven hospitalized COVID-19 patients and six healthy controls, comprising approximately 44, 721 cells . The second dataset is a human single-cell atlas of fetal gene expression [1], containing 504, 028 cells from 77 main cell types, after applying the same balanced sampling strategy as in [8]. UMAP embeddings of the two datasets are shown in Supplementary Figures 2 and 3.

Standard preprocessing steps included normalization by total counts per cell, scaling to 10,000 counts per cell, and applying a log(1 + *p*) transformation to stabilize variance. We then selected the top 2,000 highly variable genes to preserve the most biologically informative features while reducing technical noise. Principal component analysis (PCA) was then performed using the default implementation in *scikit-learn* to maintain a consistent latent space across all methods.

For all state-of-the-art methods, we used the official GitHub implementations (see Supplementary Section 4) with their default configurations unless otherwise noted. Because the original SEACells implementation was infeasible to run on the human fetal atlas dataset, we applied a set of low-level, algorithm-neutral optimizations to its Python/SciPy codebase. These modifications removed redundant Python-level loops and refactored several core routines into vectorized, sparse-efficient operations. In the original version, constructing the kernel matrix alone required more than 22 hours on the human fetal atlas dataset. After optimization, the same step completed in approximately 8 minutes, representing a speedup of over 160*×*. For MetaQ on the same large dataset, we additionally reduced the number of training epochs from default 300 to 40, decreased the convergence threshold from 10 to 2, and increased the batch size from 512 to 1024 to ensure convergence within available computational resources. All methods were executed through the integrated benchmarking pipeline of [7] to ensure consistency and reproducibility.

All computational experiments in this study were performed on a high-performance computing cluster. Each job was executed on a single node with an Intel Xeon Gold 6226R CPU (2.90 GHz) and 120 GB of RAM for both the PBMC and the human fetal atlas datasets. No GPU acceleration was used in any experiment, and each run was limited to a maximum of 48 hours.

### 3.2 Metacell quality metrics

We evaluated intrinsic structural and biological coherence of metacells. Specifically, we implemented established quality metrics that are widely applied in metacell benchmarking studies [25][13] to evaluate the homogeneity of cells within metacells and the heterogeneity across metacells. As noted in [25], comparing compactness and separation across methods can be biased when each method operates in a different latent space. To mitigate this potential unfairness, all methods were provided identical PCA-transformed data, ensuring a consistent geometric basis for comparison. For a given metacell *m* ∈ [*k*] where *k* represents the total number of metacells, we summarize the metrics in Table 1 below.

**Table 1.** Metacell Quality Metrics.

| Metric | Definition | Desirable Direction |
| --- | --- | --- |
| **Compactness** | $\text{compactness}(m) = -\frac{1}{N} \sum_{i=1}^N \text{Var}(\vec{x}_i^m)$ | ↑ (higher = more homogeneous) |
| **Separation** | $\text{separation}(m) = \min_{\ell \neq m} \text{dist}(m, \ell)$ | ↑ (higher = better separation) |
| **SC Ratio** | $\text{SC Ratio}(m) = \frac{\text{separation}(m)}{\sqrt{-\text{compactness}(m)}}$ | ↑ (higher = better balance) |
| **ECDF** | $\text{ECDF}(t) = \frac{1}{k} \sum_{i=1}^k \mathbf{1}_{\{x_i \leq t\}}$ | → (flatter / right-shifted = better) |
| **Purity** | $\text{purity}(m) = \max_j \frac{\vert M \cap C_j\vert }{\vert M\vert }$ | ↑ (higher = more coherent) |
| **INV** | $\text{INV}(m) = \text{P}_{95} \left( \frac{\text{Var}(\vec{x}_g^m)}{\text{Mean}(\vec{x}_g^m)} \right)$ | ↓ (lower = less variability) |

Compactness measures how tightly cells belonging to a metacell cluster together in the latent space, where 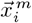 represents the vector of the *i*-th diffusion component across all cells assigned to metacell *m*, and *N* is the embedding dimensionality. Note that we negate the compactness defined in [25] without loss of generality. Separation quantifies how distinct each metacell is from its nearest neighbor in latent space, where dist(*m, ℓ*) is the Euclidean distance between the centroids of metacells *m* and *ℓ* in diffusion space.

Due to the inherent tradeoff between compactness and separation discussed in [25], we propose a combined metric, the SC Ratio, to jointly evaluate the balance between intra-metacell homogeneity and inter-metacell distinctness. To visualize the distributional characteristics of SC Ratio values, we use the empirical cumulative distribution function (ECDF), defined for a set of *k* SC Ratio values {*x*_1_, *x*_2_, …, *x*_*k*_} for *k* metacells, where 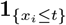 is the indicator function that equals 1 if *x*_*i*_ ≤ *t* and 0 otherwise.

Purity evaluates the consistency of celltype composition within each metacell. For a metacell *m* and ground-truth celltype labels {*C*_*j*_ }, denote *M* as the set of cells belonging to metacell *m*, and *C*_*j*_ is the set of cells annotated as label *j*. The INV metric captures the internal variability of gene expression within each metacell relative to its mean level, focusing on highly variable genes, where 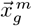 denotes the vector of expression values for gene *g* across cells in metacell *m*, and P_95_ denotes the 95th percentile computed over all genes.

Furthermore, we evaluated how well metacell representations preserve biologically meaningful signal in a downstream analysis task, namely cell-type classification. We computed balanced accuracy using the CellTypist majority voting framework [26]. Balanced accuracy accounts for potential class imbalance by averaging the recall (true positive rate) across all *T* cell types:

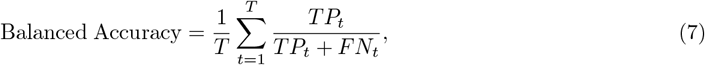

where *TP*_*t*_ and *FN*_*t*_ denote the number of true positive and false negative predictions for cell type *t*, respectively. This metric provides an unbiased estimate of classification performance even when the celltype distribution is highly imbalanced.

### 3.3 Coresets produce geometrically and biologically coherent metacells

From the top row of Figure 1, we observe that, for most methods, except MetaCell2, compactness increases monotonically with the number of metacells, indicating that finer partitions capture increasingly homogeneous transcriptional neighborhoods in the PBMC dataset. Conversely, separation generally decreases as expected, as metacell centroids move closer in higher-resolution partitions. The irregular pattern of MetaCell2 likely arises from the need to generate smaller γ values to match the same number of metacells shown on the x-axis, effectively altering its resolution and destabilizing graph partitions.

**Figure 1.**
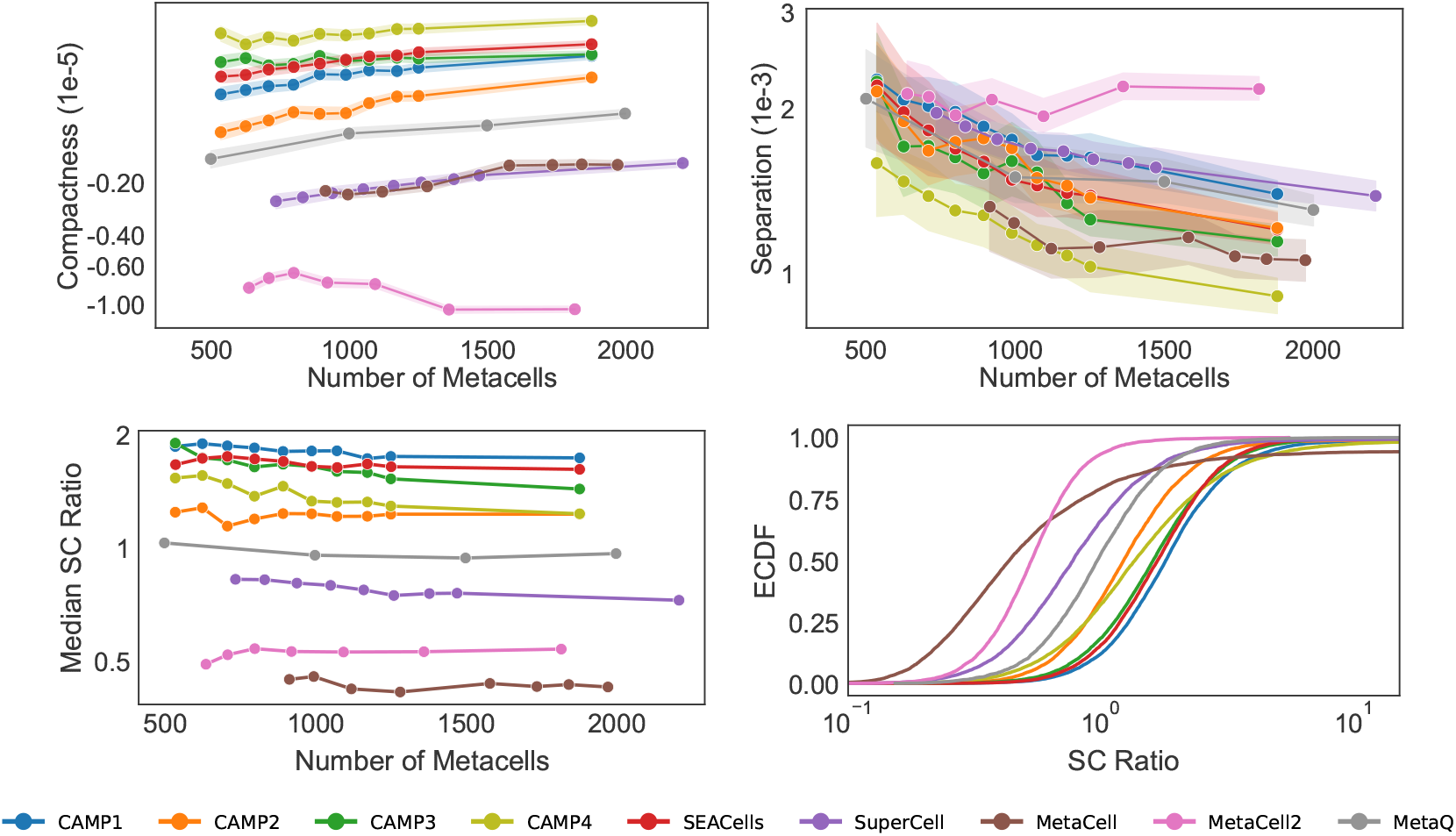
Compactness and separation of metacells produced by competing methods on the PBMC dataset. Because MetaCell and MetaCell2 do not support explicitly specifying the number of metacells, their results appear slightly shifted along the x-axis. One SuperCell point was omitted to maintain consistent resolution levels across methods.

In contrast, all CAMP variants, especially CAMP1, achieve strong compactness and separation profiles, ranking consistently among the top methods. CAMP2’s slightly lower SC ratio reflects using *O*(*k*) instead of its kernel-driven *O*(*nk*) coreset requirement derived from (4), yet it remains highly competitive (see Section 4). The ECDF plot (Figure 1, bottom right) further underscores CAMP’s stability: CAMP1 exhibits the most right-shifted distribution and CAMP4 the flattest, whereas the ECDF curve for MetaCell2 is both left-shifted and steep despite its higher separation.

For the human fetal atlas dataset, we observe the same overall trends (Figure 2). MetaQ, in contrast, suffers from unstable vector-quantization behavior on large datasets. Among all methods, CAMP1 and CAMP4 consistently yield the highest compactness and SC ratio values. SEACells achieves slightly higher separation values than CAMP1 and CAMP2, but this comes at the cost of substantially lower compactness. From the ECDF plot, all CAMP variants exhibit right-shifted and flatter distributions, while competing methods such as MetaQ and SuperCell show steeper or left-skewed curves. For the human fetal atlas dataset, the optimal γ settings [25] for MetaQ may fall outside the range evaluated here, potentially below 1, 000 metacells or above the highest resolution of 25, 000 examined in this study.

**Figure 2.**
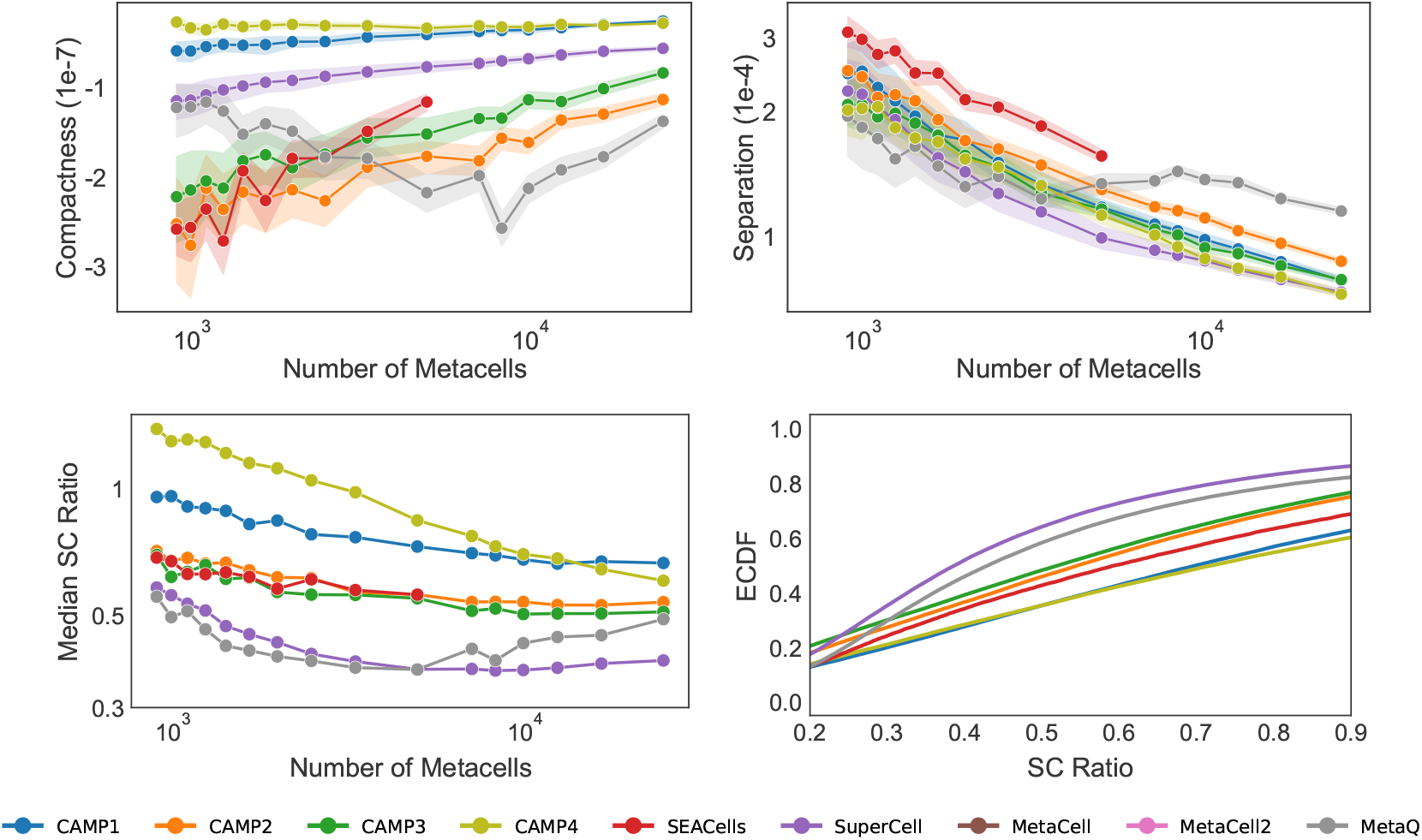
Compactness and separation of metacells returned by competing methods on the human fetal atlas dataset.

The purity trend closely parallels that of compactness, which is expected since cells with similar transcriptional profiles are more likely to belong to the same cell type. Consistent with their performances in terms of compactness, MetaCell2 in the PBMC and MetaQ in the human fetal atlas data from Figure 3 exhibit poor performance due to their smaller γ and unstable quantization, respectively. In contrast, our CAMP framework demonstrates consistent and biologically coherent performance. Among all methods, CAMP4 achieves the highest purity and lowest INV, indicating the most homogeneous and well-defined metacell compositions for both PBMC and the human fetal atlas dataset. The default variant, CAMP1, is competitive in terms of Purity across both datasets, outperforming MetaQ and MetaCell2 on the PBMC data, and MetaQ and SEACells on the human fetal atlas data.

**Figure 3.**
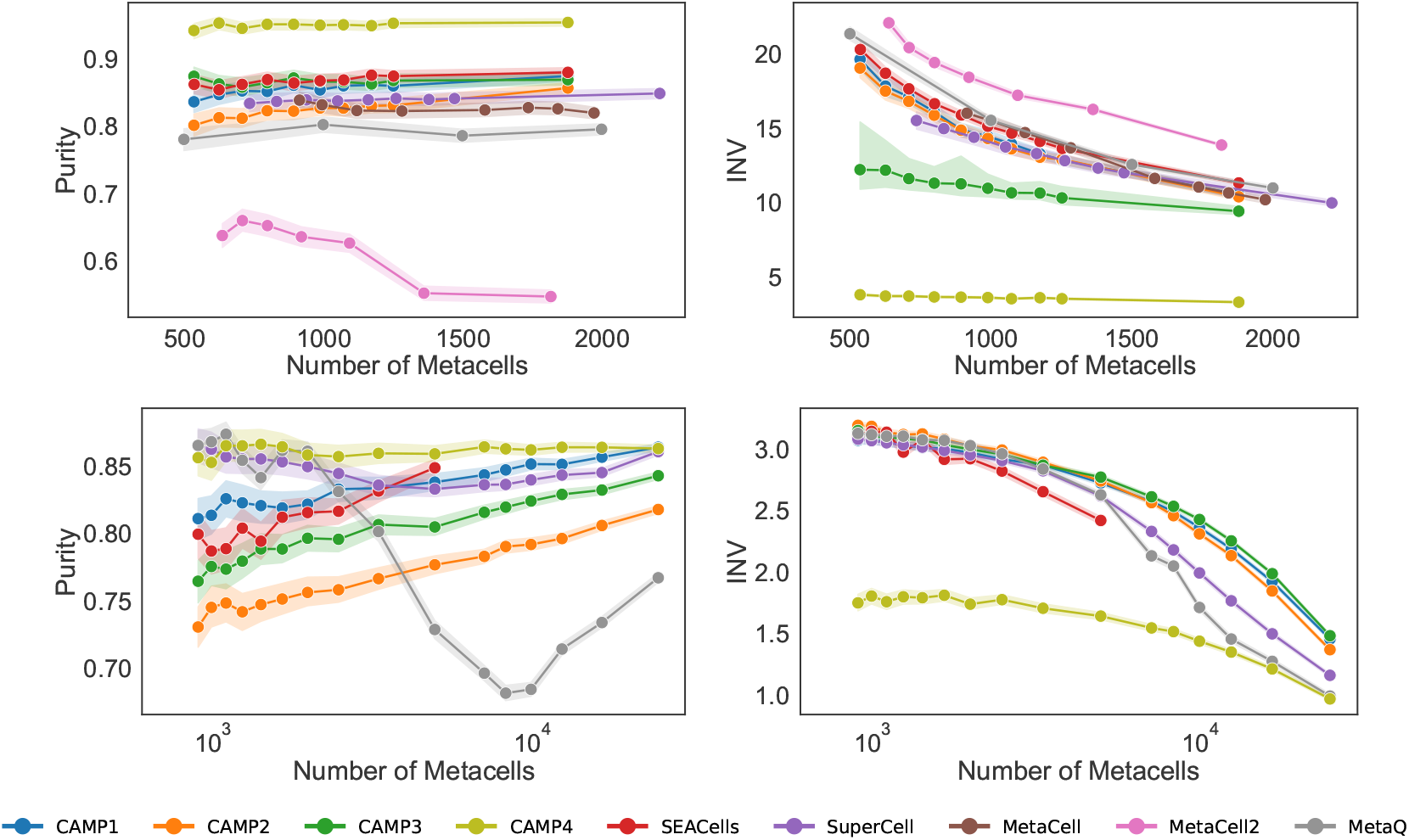
Purity and INV scores of metacells returned by competing methods on the PBMC (top) and the human fetal atlas datasets (bottom).

### 3.4 Fast and memory-efficient metacell construction

For the PBMC dataset (Figure 4, left), all CAMP variants consistently occupy the lower end of the runtime spectrum. At a resolution of 500 metacells, CAMP1 completes in approximately 1 second, whereas MetaQ requires 12,615 seconds, about four orders of magnitude slower. At finer resolutions near 2,000 metacells, CAMP1 finishes in 3 seconds while MetaQ takes 14,483 seconds, a difference of roughly 3.5 orders of magnitude. SEACells, despite achieving competitive compactness, requires 5,911 seconds at a comparable metacell resolution of 1,879, which is nearly three orders of magnitude slower than CAMP1. These discrepancies are consistent across the entire resolution range and highlight the substantial cost of iterative kernel-based updates in SEACells and the autoencoder training steps of MetaQ.

**Figure 4.**
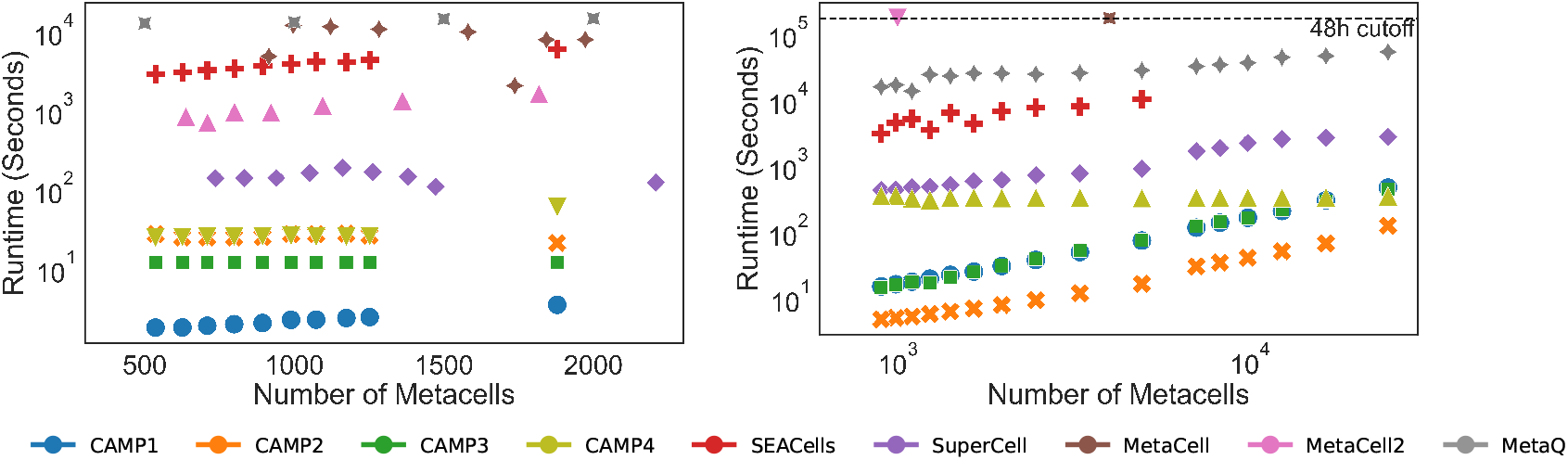
Runtime in seconds on the PBMC (left) and the human fetal atlas (right) datasets.

For the human fetal atlas dataset (Figure 4, right), all CAMP variants completed partitioning of the 504,028 cells in under 8 minutes across all metacell scales, with CAMP1-3 computing distances and similarities on-the-fly. In contrast, MetaCell and MetaCell2 each reached the 48-hour runtime ceiling, while SEACells required 10,189 seconds, which is more than an order of magnitude slower than CAMP, and ultimately failed with out-of-memory errors at higher resolutions approaching 7,000 metacells. MetaQ also becomes substantially slower at large scales, requiring 52,971 seconds at 25,000 metacells, approximately 113*×* slower than CAMP (nearly two orders of magnitude). SuperCell performs moderately faster than other non-CAMP methods due to its approximation parameter (see Supplementary Section 4), yet it still remains slower than all CAMP variants.

### 3.5 Accurate downstream classification from coreset-based metacells

In this section, we examine whether coreset-based partitions preserve sufficient biological signal to support accurate downstream classification. We assessed cell type prediction performance using CellTypist [26] with majority voting, reporting the balanced accuracy across the ten largest cell-type classes in each dataset. As no method reliably recovered the remaining low-frequency cell types, these minority populations were omitted from the quantitative analysis.

For the PBMC dataset, we assessed post hoc performance at a resolution of 1,000 metacells (Figure 5), as this is the common resolution at which all methods produce results. For the human fetal atlas data, results corresponding to 3,000 metacells are shown in Figure 6. Additional results with other metacell resolutions are provided in Supplementary Figures 4–9. The y-axis denotes the true labels and the x-axis the predicted labels, both sorted in descending order by class size (top to bottom and left to right). The corresponding cell types are listed in Supplementary Section 5.

**Figure 5.**
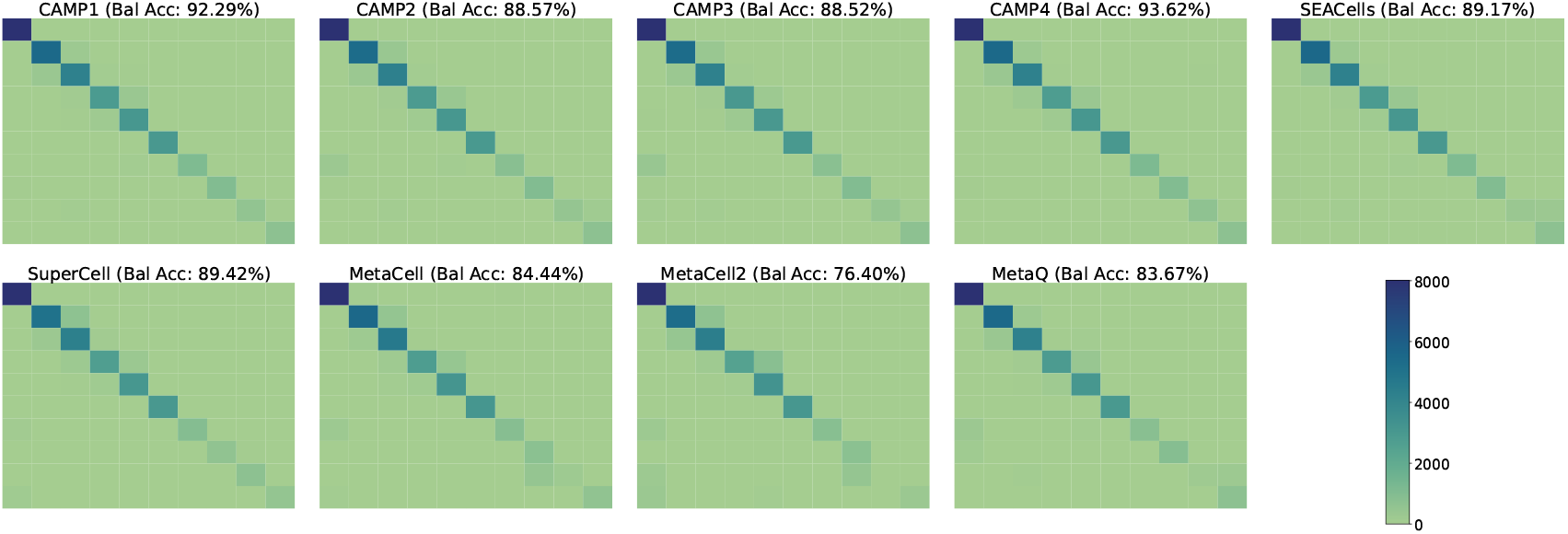
Cell-type confusion heatmaps for the ten largest cell-type classes based on 1,000 metacells computed by competing methods on the PBMC dataset. Balanced accuracy (Bal Acc) is reported on the top.

**Figure 6.**
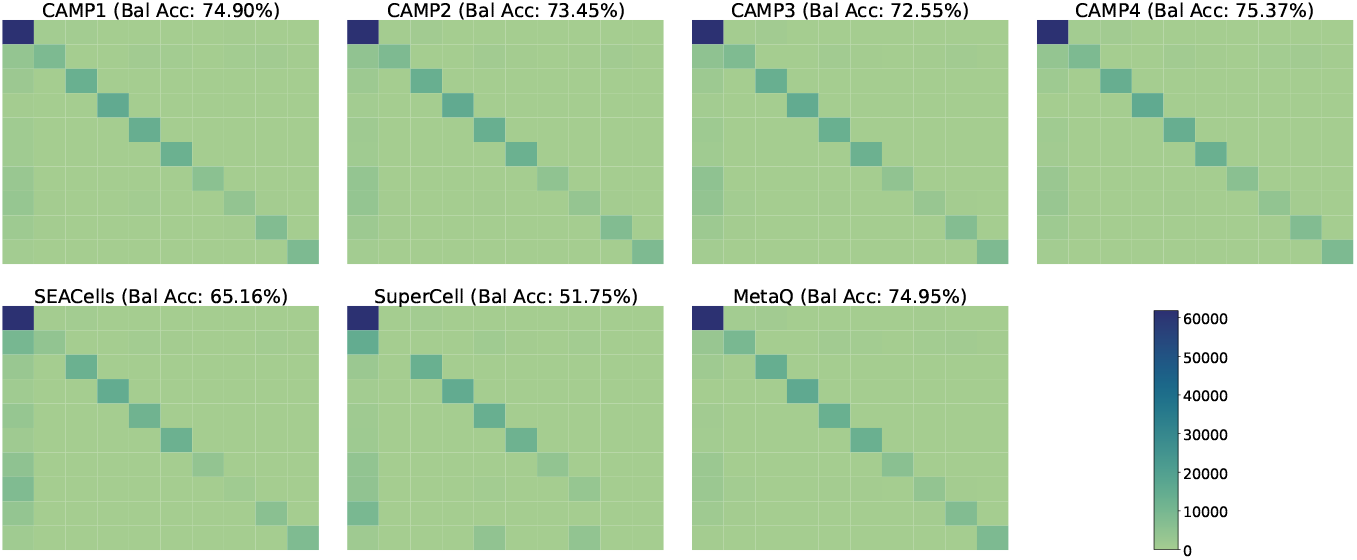
Cell-type confusion heatmaps for the ten largest cell-type classes based on 3,000 metacells computed by competing methods on the human fetal atlas dataset. Balanced accuracy (Bal Acc) is reported on the top.

Across both datasets, CAMP variants achieved the strongest overall performance. On the PBMC dataset (Figure 5), CAMP1 and CAMP4 obtained the highest balanced accuracies of 92.29% and 93.62%, respectively. In contrast, SEACells and MetaQ frequently misclassify Class-switched B cells as IgG plasmablasts, while MetaCell and MetaCell2 tend to confuse them with CD16 monocytes. SuperCell and MetaCell2 also show notable confusion between NK and CD8m T cells. On the human fetal atlas dataset (Figure 6), CAMP1, CAMP4, and MetaQ similarly achieved the top balanced accuracies (74.90%, 75.37%, and 74.95%), maintaining clean diagonal patterns in their confusion matrices. Most competing methods, except CAMP4 and MetaQ, show substantial confusion among neural cell types, especially assigning Inhibitory or Limbic neurons to the Excitatory neuron class. SEACells also performs poorly on Ganglion cells, often grouping them together with Excitatory neurons.

## 4 Conclusion

In this work, we introduced CAMP, a scalable and geometry-preserving framework for metacell construction that unifies coreset theory with archetypal analysis to address the computational and biological challenges of large-scale single-cell transcriptomics. By leveraging lightweight *k*-means coresets, CAMP identifies a small but highly representative subset of cells that captures the geometry of the transcriptomic manifold with high probability. This design provides strong theoretical guarantees while reducing both runtime and memory usage by orders of magnitude compared to existing metacell frameworks, including MetaCell, MetaCell2, SuperCell, SEACells, and MetaQ.

In our experiments, CAMP consistently produced compact, well-separated, and biologically coherent metacells. These geometric advantages translate into practical downstream benefits: CAMP1 and CAMP4 achieved the highest balanced accuracies in cell-type classification, demonstrating that CAMP preserves biologically meaningful variation even under aggressive compression. Importantly, CAMP attains this accuracy without incurring the heavy computational demands characteristic of prior methods. All CAMP variants completed metacell construction from half a million cells in under 8 minutes on CPU-only hardware, whereas several state-of-the-art approaches either exceed the 48-hour runtime limit or fail due to memory constraints. This establishes CAMP as a uniquely efficient and robust paradigm for atlas-scale metacell inference.

Collectively, its four variants allow CAMP to adapt to a wide range of biological settings. As summarized in Table 2 below, CAMP1 serves as a fast and reliable default; CAMP2 provides the most scalable option through fully vectorized linear-kernel updates; CAMP3 captures nonlinear and density-dependent structure when additional flexibility is required; and CAMP4 offers a balanced hybrid, trading a small amount of additional computation for greater robustness across heterogeneous transcriptomic landscapes. On the theoretical side, CAMP clarifies the relationship between coreset size and metacell complexity. The required sampling rate follows the bounds derived in Section 2.1: *m* = *O*(*k*) for CAMP1, *m* = *O*(*nk*) for CAMP2, and *m* = *O*(*k*^2^) for CAMP3. Although CAMP2 and CAMP3 have larger theoretical requirements due to their kernel constructions, our empirical findings demonstrate that sampling only *m* = *O*(*k*) points is sufficient across diverse biological settings. This observation highlights an intriguing gap between worst-case analysis and practical behavior, and motivates future work on tightening theoretical guarantees for kernel-based CAMP variants.

**Table 2.** CAMP variant selection guide across benchmark metrics. Streaming refers to efficient updates computed on-the-fly, i.e., only when needed, without constructing the full kernel or distance matrix in advance.

| Variant | Kernel | Compactness | Separation | Purity | INV | Accuracy | Streaming |
| --- | --- | --- | --- | --- | --- | --- | --- |
| **CAMP1** | Default Free | ✓ | ✓ | ✓ |  | ✓ |  |
| **CAMP2** | Linear |  | ✓ |  |  |  | ✓<sup>1</sup> |
| **CAMP3** | Non-linear | ✓ |  | ✓ |  |  |  |
| **CAMP4** | Hybrid | ✓ |  | ✓ | ✓ | ✓ |  |

CAMP also raises several conceptual questions for methodological theory. Understanding how geometric summarization interacts with downstream tasks, including differential expression, clustering stability, and rare-cell detection. This could inspire metacell constructions tailored to specific biological objectives. Looking ahead, CAMP provides a modular foundation for several promising extensions. Geometry-preserving coresets could accelerate multi-omic metacell construction for ATAC-seq, CITE-seq, methylation, or spatial datasets, and incorporating trajectory-aware or pseudotime-informed constraints may allow CAMP to better capture continuous developmental processes.

All datasets, together with the complete CAMP implementation pipeline and reproducible scripts for all competing methods (including our SEACells optimizations), are publicly available in our GitHub repository: https://github.com/danrongLi/CAMP.

## Footnotes

1 Although CAMP1–3 support on-the-fly (streaming) updates, only CAMP2 enables computationally efficient streaming. Its linear kernel allows fully vectorized operations in Python, enabling fast, loop-free batched dot-product updates.

