## Supplementary for "CAMP: Coreset Accelerated Metacell Partitioning enables scalable analysis of single-cell data"

#### 1 Seed distribution for CAMP1-3

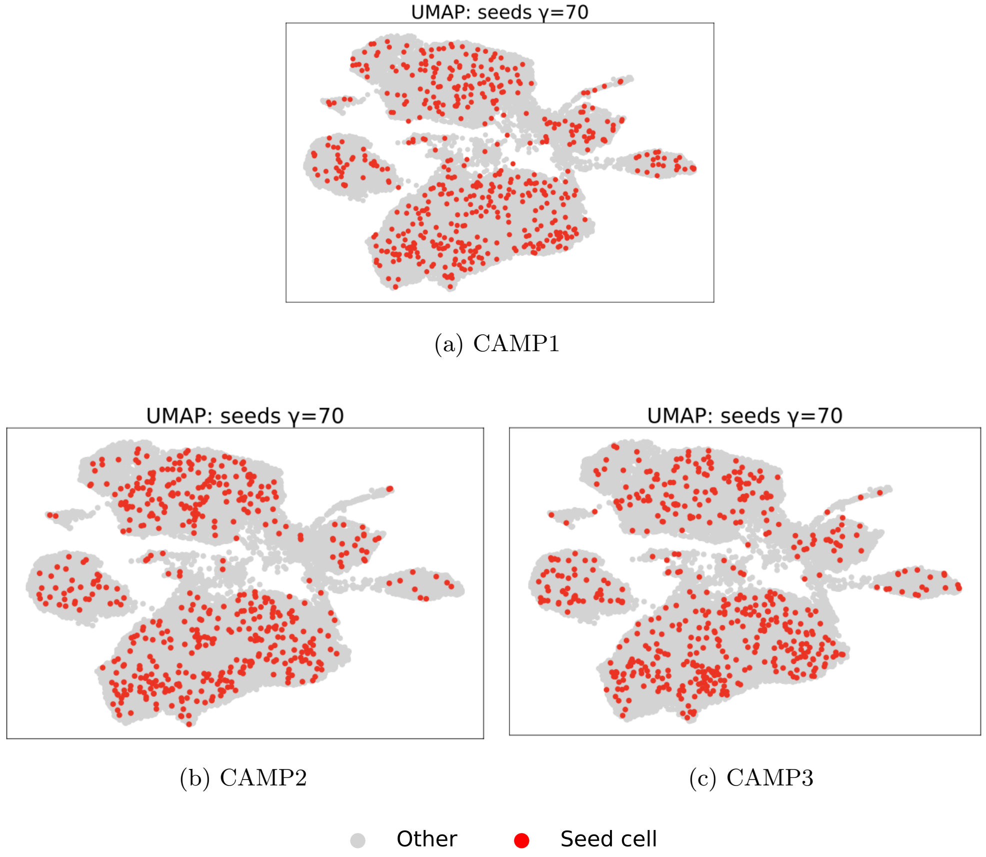


(a) CAMP1

(b) CAMP2

(c) CAMP3

Figure 1: Seed distribution on the PBMC dataset.

Figure 1 shows that even when omitting a computationally expensive archetypal analysis, the lightweight coresest accurately covers the the inner region.

*Corresponding authors

#### 2 Refinement Algorithm

**Algorithm 1 CAMP Refinement (Lloyd-style updates for CAMP1-3)**

**Require:** Dataset $\mathcal{X}$, initial assignments $z(x)$ and centers $\{\mu_j\}_{j=1}^m$ from initial CAMP step in Algorithm 1
**Require:** Maximum iterations $T \leftarrow 10$
**Ensure:** Refined membership map $z : \mathcal{X} \to \{1, \ldots, m\}$

**for** $t = 1$ to $T$ **do**
  **for** $j = 1$ to $m$ **do**
    $S_j \leftarrow \{x \in \mathcal{X} : z(x) = j\}$
    **if** $|S_j| > 0$ **then**
      $\mu_j \leftarrow \frac{1}{|S_j|} \sum_{x \in S_j} x$ ▷ update center to cluster mean
    **else**
      $x^\star \leftarrow \arg\max_{x \in \mathcal{X}} \min_{\ell \in \{1,\ldots,m\}} d(x, \mu_\ell)^2$ ▷ farthest unassigned point
      $\mu_j \leftarrow x^\star$ ▷ re-seed empty cluster
    **end if**
  **end for**
  **for** $x \in \mathcal{X}$ **do**
    $z(x) \leftarrow \arg\min_{j \in \{1,\ldots,m\}} d(x, \mu_j)$ ▷ reassign to nearest updated center
  **end for**
**end for**
**return** $z$

#### 3 UMAP embeddings of PBMC and human fetal atlas data

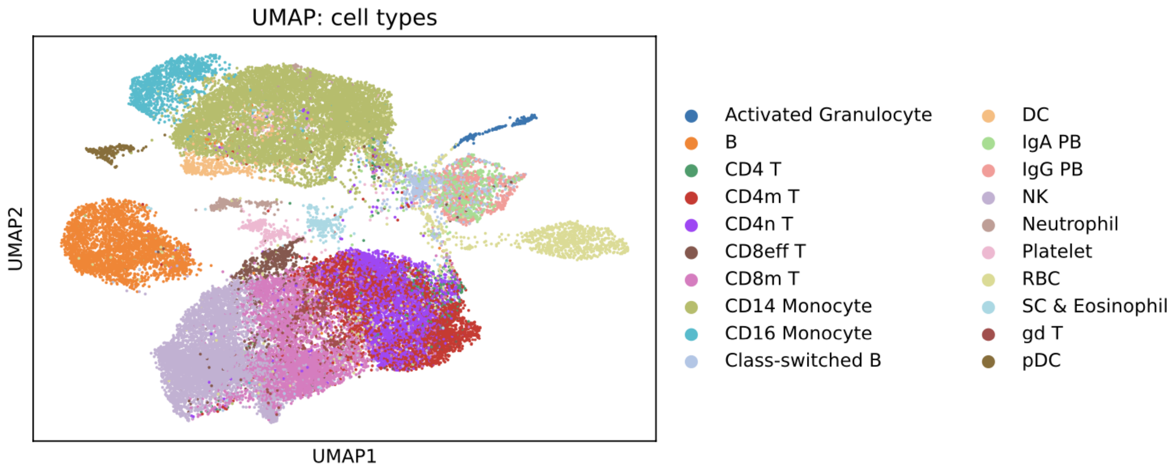


Figure 2: UMAP embedding of the PBMC dataset.

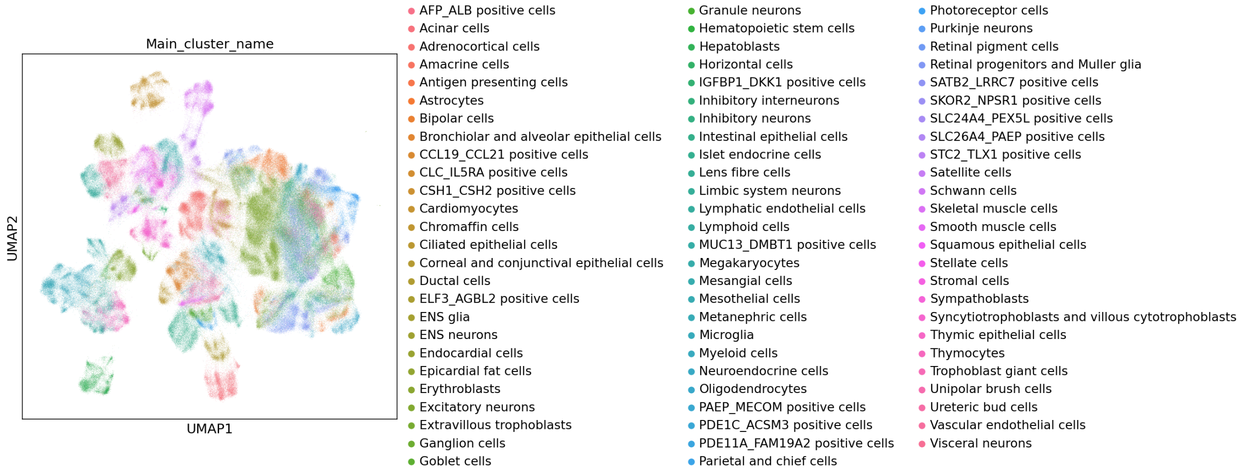


Figure 3: UMAP embedding of the human fetal atlas dataset.

### 4 Software packages used for state-of-the-art methods

For SEACells, we optimized it by modifying `cpu.py` and `build_graph.py` files in the original Python implementation (`https://github.com/dpeerlab/SEACells`) and used this optimized version with `use_sparse=True`, which is available only in CPU mode. SEACells produced out-of-memory errors on the human fetal atlas dataset whenever the target average number of cells per metacell, $\gamma$, was $\leq 70$. For SuperCell, we used the R package (`https://github.com/GfellerLab/SuperCell`) with the option `do_approx=True`. For MetaQ, we used the Python implementation (`https://github.com/XLearning-SCU/MetaQ`) with default parameters on the PBMC dataset. On the human fetal atlas dataset, we changed the default parameters as discussed in main text. MetaCell and MetaCell2 were run using the R package (`https://tanaylab.github.io/metacell/`) and the Python package (`https://pypi.org/project/metacells/`), respectively. On the human fetal atlas dataset, MetaCell (configured with `knn = 50` and `amp = 1`) exceeded the 48-hour runtime limit, and MetaCell2 exceeded the limit for $\gamma = 900$ and $\gamma = 500$.

### 5 Ten largest cell-type classes

| PBMC | human fetal atlas |
|---|---|
| CD14 Monocyte | Excitatory neurons |
| NK | Inhibitory neurons |
| CD8m T | Astrocytes |
| CD4m T | Adrenocortical cells |
| CD4n T | Granule neurons |
| B | Purkinje neurons |
| RBC | Metanephric cells |
| CD16 Monocyte | Ganglion cells |
| Class-switched B | Limbic system neurons |
| IgG PB | Intestinal epithelial cells |

Table 1: Top 10 cell types in terms of class size in descending order from top to bottom for both the PBMC and the human fetal atlas datasets.

### 6 Cell type prediction on the human fetal atlas dataset

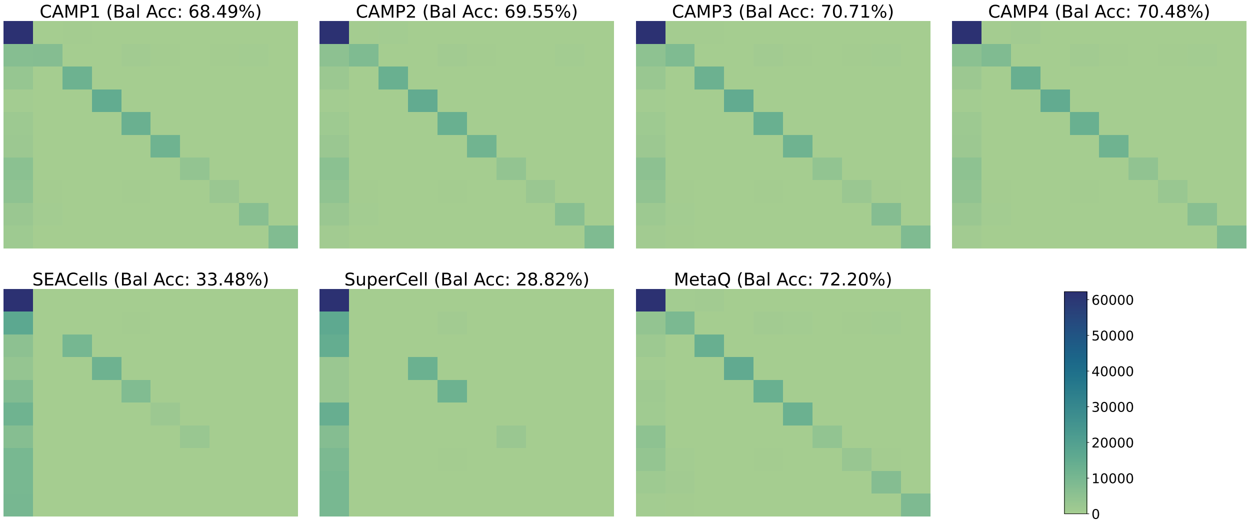


Figure 4: Cell-type confusion heatmaps for the ten largest cell-type classes based on 1,000 metacells computed by competing methods on the human fetal atlas dataset. Balanced accuracy (Bal Acc) is reported on the top.

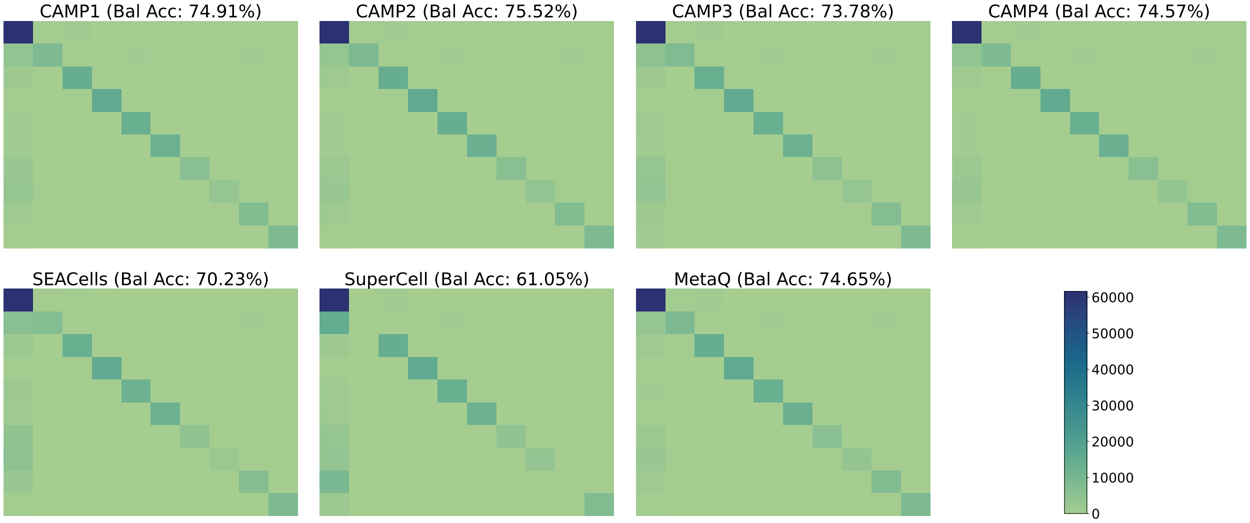


Figure 5: Cell-type confusion heatmaps for the ten largest cell-type classes based on 5,000 metacells computed by competing methods on the human fetal atlas dataset. Balanced accuracy (Bal Acc) is reported on the top.

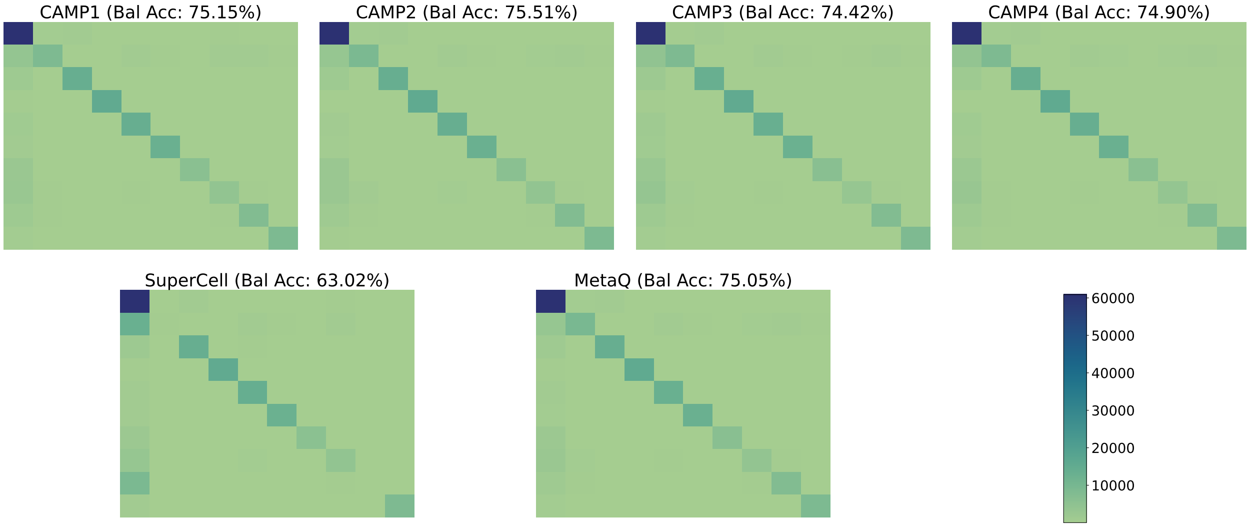


Figure 6: Cell-type confusion heatmaps for the ten largest cell-type classes based on 7,000 metacells computed by competing methods on the human fetal atlas dataset. Balanced accuracy (Bal Acc) is reported on the top.

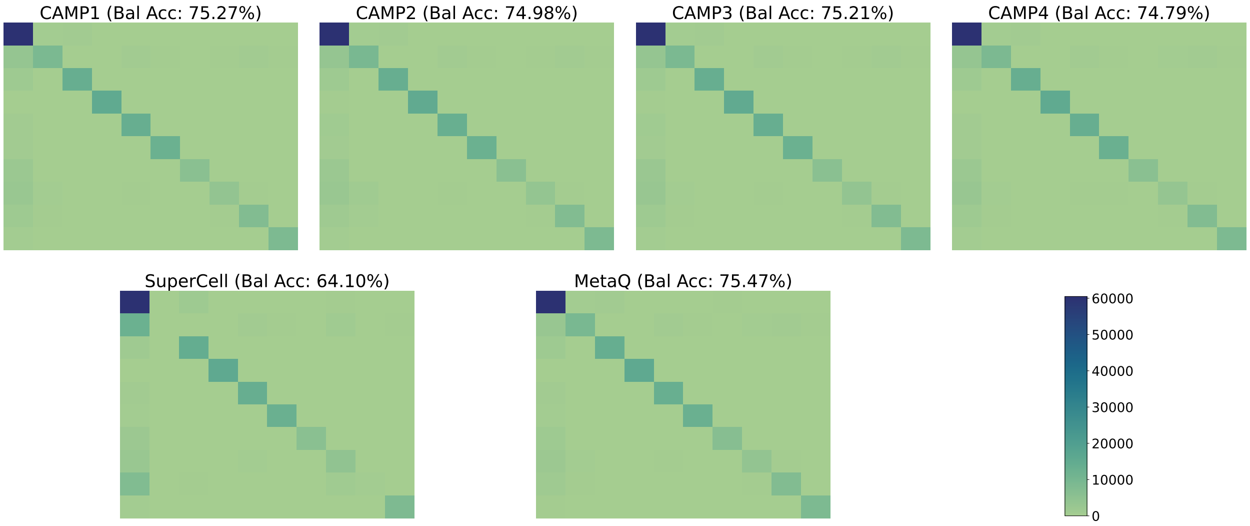


Figure 7: Cell-type confusion heatmaps for the ten largest cell-type classes based on 10,000 metacells computed by competing methods on the human fetal atlas dataset. Balanced accuracy (Bal Acc) is reported on the top.

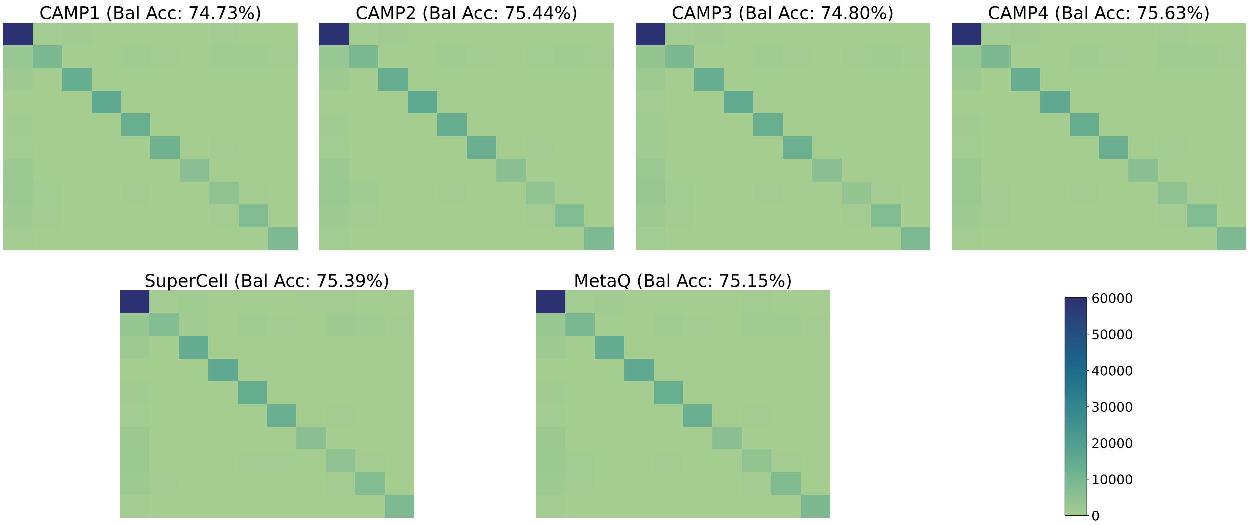


Figure 8: Cell-type confusion heatmaps for the ten largest cell-type classes based on 15,000 metacells computed by competing methods on the human fetal atlas dataset. Balanced accuracy (Bal Acc) is reported on the top.

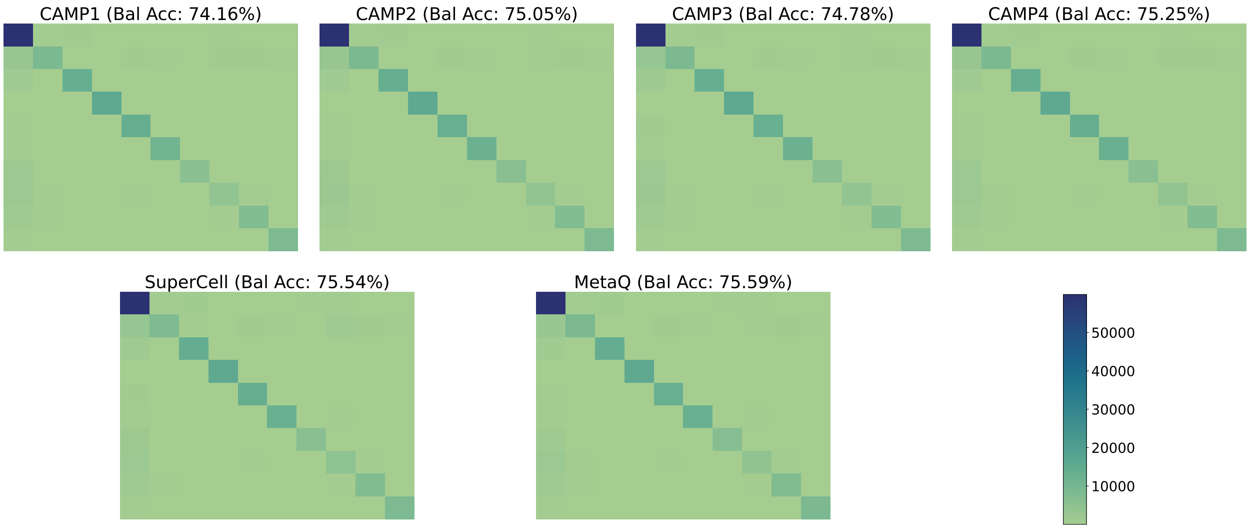


Figure 9: Cell-type confusion heatmaps for the ten largest cell-type classes based on 25,000 metacells computed by competing methods on the human fetal atlas dataset. Balanced accuracy (Bal Acc) is reported on the top.
